# SARS-CoV E protein couples asymmetric leaflet thickness and curvature deformations

**DOI:** 10.1101/2025.04.18.649534

**Authors:** Jesse W Sandberg, Grace Brannigan

## Abstract

The Envelope protein (E protein) of SARS-CoVs 1 and 2 has been implicated in the viral budding process and maintaining the spherical shape of the virus, but direct evidence linking the protein to long-range membrane deformation is still lacking. Computational predictions from molecular simulation have offered conflicting results, some showing long-range E-induced membrane curvature and others showing only local deformations. In the present study, we determine the mechanism driving these deformations by modulating the degree of hydrophobic mismatch between protein and membrane. We observe that certain barostat and restraint settings, common in coarse-grained MD simulations, can prevent equilibration of the membrane area. Our results indicate that the E protein does not induce long-range curvature, but does exhibit severe local deformations that are exacerbated by hydrophobic mismatch. These deformations occur in conjunction with local leaflet thickness asymmetry, suggesting asymmetry and curvature couple to reduce the free energy cost of a deformed membrane.

**Highlights:**

- E protein pentamers from SARS-CoV-1 and SARS-CoV-2 do not induce long-range curvature in membranes when simulated in isolation.
- Previous findings documenting long-range membrane deformation may reflect the use of restraint and barostat settings that trap the membrane in a compressed state.
- E proteins induce large, asymmetric membrane deformations local to the protein, but these deformations do not propagate into the bulk.
- Membrane leaflet thickness asymmetry may be non-negligible around proteins that are not cylindrical.
- Membrane leaflet thickness asymmetry and mean curvature couple to alleviate free energy cost of deformation.

**Graphical Abstract:** 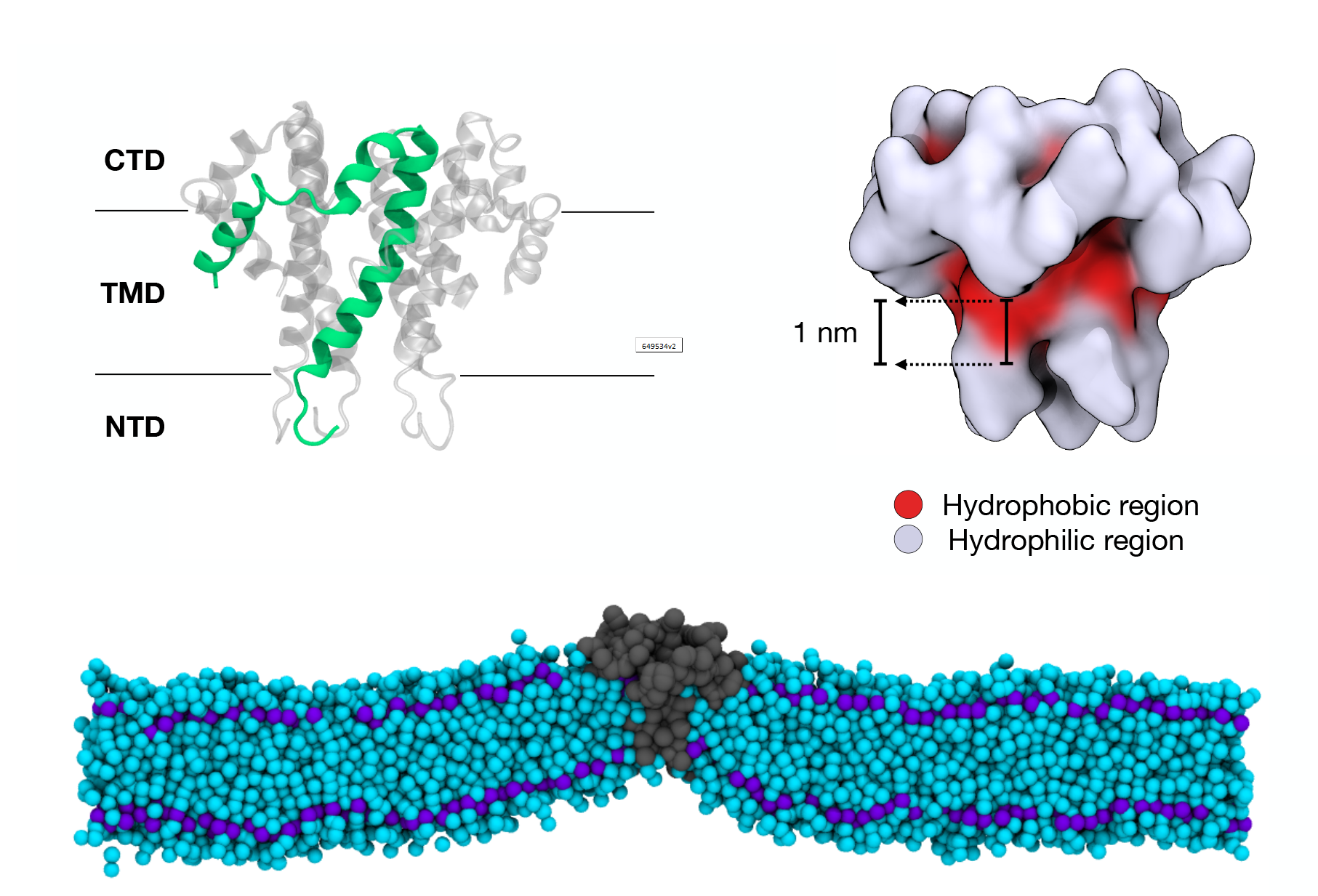

## 1. Introduction

The Envelope protein (E protein) is a structural protein present across many members of the Coronaviridae virus family [1]. Although its precise functions remain undetermined, knockout or mutation of E proteins in several coronaviruses (CoVs) has led to reduced viral titres and reproductive ability [2–8], suggesting a critical role for the protein in the viral life cycle. While the extent of genetic conservation varies across family members, sequence similarity is high enough among human CoVs (HCoVs) that the E protein has been suggested as a target for pan-HCoV screening products [9]. The protein sequence is most conserved across those HCoVs that cause Severe Acute Respiratory Syndrome (SARS) [10–12]. Over the course of the SARS-CoV-2 pandemic, the E protein accumulated the fewest mutations out of any of the four structural proteins [9, 13–17] further highlighting that the E protein remains a stable target for therapeutic intervention as coronaviruses continue to evolve.

Electrophysiological studies show that the E protein is a viroporin, although there is some debate about the selectivity and environmental dependence of the channel [18–28]. The ion channel activates the host cell inflammasome via Ca^2+^ transport [29–32], and de-acidifies lysosomes to allow for viral egress from an infected cell [33]. While the E protein is only present in low copy number in the viral envelope, it is highly expressed in the ERGIC of an infected cell [23, 34, 35], where it plays a significant role in the viral assembly and budding process [1, 36–38]. Schoeman and Fielding [1] speculate that the E protein contributes to the budding process by inducing membrane curvature.

The E protein is the smallest of the four SARS-CoV structural proteins. Its sequence contains a 7-residue N-terminal domain (NTD), a 31-residue transmembrane domain (TMD), and a 37-residue C-terminal domain (CTD) (Fig. 1 A&B). It has been shown to oligomerize into a homo-pentamer [25, 32, 39–41], but other forms also appear to be possible [42, 43]. Structures of the E protein determined in micelles have shown the CTD to be composed primarily of alpha helices [25, 44], although another group has found that E proteins expressed in a bilayer environment adopt a *β*-sheet CTD conformation [45, 46]. Some have therefore conjectured that the protein may shift its conformation depending on the curvature of the environment, but to our knowledge this linkage has never been demonstrated. Furthermore, the *β*-sheets in Ref.s [45, 46] were only found at protein-to-lipid ratios so high as to suggest protein-protein aggregation. The only pentameric structure available from the Protein Data Bank with the CTD present [25] (depicted in Fig. 1) shows the CTD descending toward the NTD, potentially occluding half of the TMD helix from contact with the membrane. We elected to use this structure and a protein-to-lipid ratio far lower than that used in Ref.s [45, 46] for our study.

**Figure 1.**
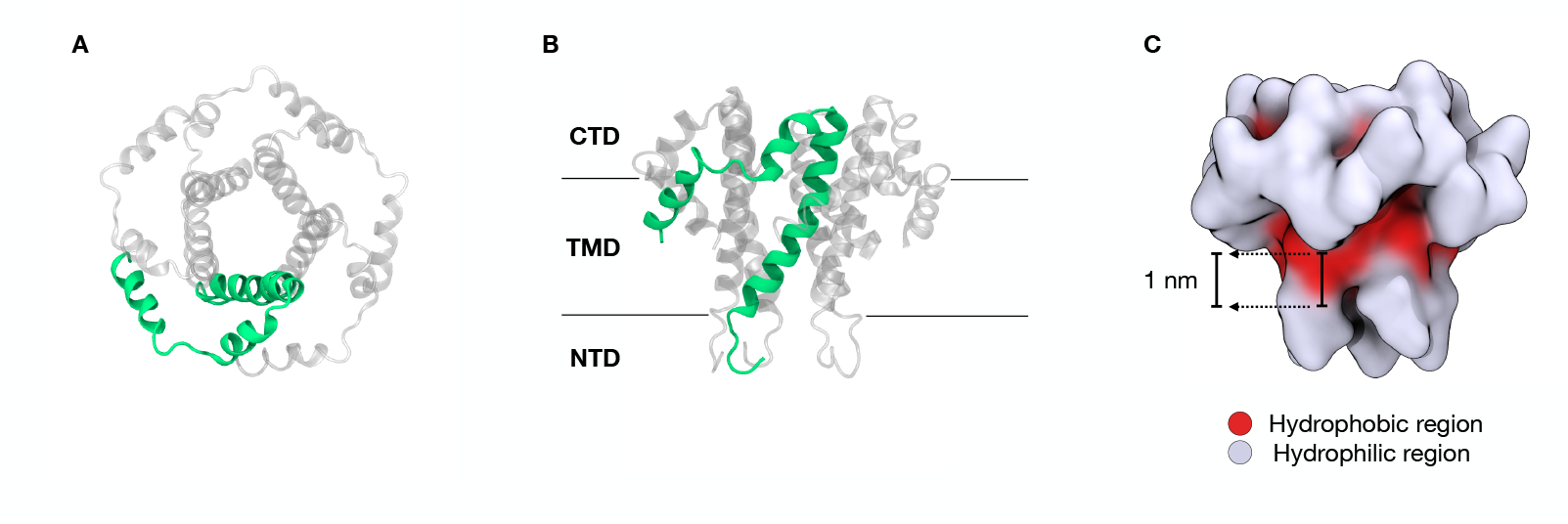
E protein structure and hydrophobic region. ‘New cartoon’ depiction of E protein pentamer (PDB ID: 5X29) as viewed from the CTD side (A) and membrane side (B). One subunit of the pentamer is colored green to highlight the structure and symmetry. Black bars in *B* indicate a naive estimate of membrane position. C) Actual hydrophobic surface (red, V17 to A36) and hydrophilic surface (light purple) of the E protein. The approximate thickness of the hydrophobic region accessible to the membrane is 1 nm. Estimates derived from contiguous hydrophobicity measurements using the *blobulator* webtool [47] and measured in VMD, as described in section 2.3.1.

The irregular shape and TMD occlusion present in this structure could be responsible for inducing membrane deformations consistent with a role in budding. While to our knowledge this linkage has never been established experimentally, several computational groups have observed membranes deforming around the E protein in Molecular Dynamics (MD) simulations with results varying. Mehregan et al. [48] simulated both pentameric [25] and monomeric [44] structures of the E protein and showed that they both induced curvature. They also showed that curvature induced by the pentamer was remarkably long-range, extending to the edge of their simulation system. In contrast, Collins et al. [49] and Kuzmin et al. [50] both concluded that the E protein pentamer could not induce long-range membrane deformation on its own, although it may be attracted to regions of high curvature [50]. These groups all used different combinations of E protein structure, oligomerization status, simulation resolution, and simulation run-time settings, making comparison of results difficult.

Ultimately there has been no reconciliation between these results to date, nor a specific mechanism behind the long-range induction of curvature identified. There are three commonly discussed mechanisms through which proteins or other rigid inclusions could deform a membrane or its constituent leaflets: the scaffold mechanism, the contact-angle mechanism, and the hydrophobic mismatch mechanism. The scaffold mechanism involves the unidirectional exertion of force over a large membrane area. This is usually accomplished by large protein complexes such as ESCRT or COPII [51, 52] (and sources therein). The remaining mechanisms are specific to integral membrane proteins. The contact-angle mechanism induces conditions under which a curved membrane has a lower free energy cost than when the membrane remains flat, typically owing to the insertion of a wedge-shaped protein or inclusion [52–55]. The hydrophobic mismatch mechanism typically involves the symmetric thickness deformation of both inner and outer membrane leaflets around a protein hydrophobic surface that differs in size from the equilibrium thickness of the membrane, minimizing the curvature of the membrane in the process [56–62].

As the E protein is a small, integral membrane protein, the scaffold mechanism seems unlikely as a candidate in this case. While the E protein is not conical, its CTD is wider than the TMD which suggests the contact angle mechanism could be responsible for any E-induced membrane deformation. Given that hydrophobic mismatch is considered to deform individual leaflets symmetrically, it is unclear how mismatch could lead to a long-range induction of curvature in one direction consistent with a role in the budding process. To our knowledge, hydrophobic mismatch has never been evaluated in proteins that are not cylindrical, so the shape of the E protein may play an unexpected role in exacerbating the membrane deformation observed. In particular, we note that Watson et al. have derived an expression for membrane free energy that includes a leaflet thickness asymmetry term, *ε*, that is coupled to membrane mean curvature [63] (eqs. 18-20). It may be that the irregular shape of the E protein creates leaflet thickness asymmetry which favorably couples with membrane curvature to lower the free energy cost of a curved membrane. In this study, we set out to determine whether the pentameric structure of the E protein produces long-range membrane curvature consistent with earlier findings [48]. We also sought to determine whether those deformations are consistent with one of the established bending mechanisms available to integral membrane proteins. Accordingly, we simulated the E protein embedded in homogeneous membranes of varying thickness and flexibility to understand what role (if any) hydrophobic mismatch may be playing in the membrane deformations. We also characterized leaflet thickness asymmetry directly, as well as its relationship to mean curvature.

## 2. Methods

### 2.1. Setup

We coarse-grained the SARS-CoV E protein pentamer (PDB ID: 5X29) [25] using the *Martinize* tool [64]. We then embedded this structure in either 100% POPC (see section 2.2.2), or 9 different homogeneous membranes of varying equilibrium thickness and lipid saturation (see section 2.2.3, Table 1) using *Insane* [65]. These systems were all solvated with water and 15% NaCl. The initial box size was 40 × 40 × 25 nm^3^, allowing for the placement of at least 2,600 lipids in each leaflet and making the protein-to-lipid ratio less than 1:5000. We refer to the membrane leaflet containing the CTD as the cytoplasmic leaflet and the leaflet containing the NTD as the luminal leaflet, in accordance with the findings in Ref. [66].

**Table 1:** Membrane composition, leaflet thickness, and relative mismatch for simulation systems. Nine homogeneous membranes were simulated as part of this study, each containing 100% of a single lipid species. All lipids had PC headgroups while the acyl chain length and saturation were varied. Relative mismatch Δ*t/t*_0_ ≡ (*t*_TMD_ −*t*_0_)*/t*_0_ was determined using *t*_TMD_ = 0.5 nm and the values of *t*_0_ below. Values for *t*_0_ were determined by measuring the average leaflet thickness of a membrane with no protein embedded over the second half of a 2 *µs* trajectory.

| Martini lipid name | Carbons per tail | Tail saturation | $t_0$ (nm) | $\Delta t/t_0$ |
| --- | --- | --- | --- | --- |
| DTPC | 8 | saturated | 0.53 | -6% |
| DYPC | 12 | mono-unsaturated | 0.81 | -38% |
| DLPC | 12 | saturated | 0.89 | -44% |
| DOPC | 16 | mono-unsaturated | 1.08 | -54% |
| DPPC | 16 | saturated | 1.23 | -59% |
| DGPC | 20 | mono-unsaturated | 1.40 | -64% |
| DBPC | 20 | saturated | 1.57 | -68% |
| Custom lipid | 24 | mono-unsaturated | 1.67 | -70% |
| DXPC | 24 | saturated | 1.90 | -74% |

**Table 2:** Symbols and Notation for quantities referenced in this article.

| Symbol | Definition |
| --- | --- |
| <u>Generic symbols</u> |  |
| $L$ | The length of the simulation box in $x$ or $y$ |
| $N_M$ | The number of lipid molecules in the membrane |
| $\Sigma$ | Instantaneous area per lipid of simulation system; $\Sigma \equiv 2L^2/N_M$ |
| $r$ | Radial distance in $xy$ plane from protein center † |
| $t_{\text{TMD}}$ | Half-thickness of protein hydrophobic region † |
| $t_0$ | Equilibrium leaflet thickness of lipid species |
| $\Delta t$ | Extent of hydrophobic mismatch; $\Delta t \equiv t_{\text{TMD}} - t_0$ |
| <u>Fields calculated from simulation</u> |  |
| $t^{(1,2)}$ | Thickness of cytoplasmic ($t^{(1)}$) or luminal ($t^{(2)}$) leaflet † |
| $z^{(1,2)}$ | Height of leaflet hydrophobic surface, normal to $xy$ plane † |
| $z^+$ | Height of bilayer midplane; $z^+ \equiv (z^{(1)} + z^{(2)})/2$ † |
| $z^{(m)}$ | Height of leaflet interface; $z^{(m)} \equiv z^{(1)} - t^{(1)}$ † |
| $H^{(1,2)}$ | Mean curvature of leaflet; $H^{(1,2)} \equiv -\nabla^2 z^{(1,2)}$ |
| $H^+$ | Bilayer mean curvature; $H^+ \equiv -\nabla^2 z^+$ |
| $\varepsilon$ | Leaflet thickness asymmetry; $\varepsilon \equiv z^{(m)} - z^+ = (t^{(2)} - t^{(1)})/2$ † |
| $\tilde{t}$ | Fractional membrane thickness; $\tilde{t} \equiv (t^{(1)} + t^{(2)})/2t_0$ |
| <u>Boundary definition and values</u> |  |
| $R^{(1,2)}$ | Radius of protein in cytoplasmic ($R^{(1)}$) or luminal ($R^{(2)}$) leaflet † |
| $\tilde{t}_R$ | The value of $\tilde{t}$ at $r = R^{(1)}$ |
| $H_R^{(1,2)}$ | The value of $H^{(1,2)}$ at $r = R^{(1)}$ or $r = R^{(2)}$ |
| $H_R^+$ | The value of $H^+$ at $r = R^{(1)}$ |
| $\varepsilon_R$ | The value of $\varepsilon$ at $r = R^{(1)}$ |

† A graphical definition is provided in Figure 2.

A mono-unsaturated lipid with 6 beads per tail was not available in the standard Martini library at the time, so we constructed a custom lipid with the *Martini lipid* .*itp generator* version 6 [65] and input string

~~~
lipid-martini-itp-v06.py -alhead “C P” -allink “G G” -altail “CCDCCC CCDCCC” -alname DZPC -o DZPC.itp
~~~

This custom lipid mimicked the structure of saturated Martini lipid DXPC, but contained an unsaturated ‘D’ bead as the third bead in each tail, counting from the glycerol backbone. Bonded parameters are generated automatically and can be found in the .itp file, which is included in the *SI*.

### 2.2. Simulation details

#### 2.2.1. Common Settings

Unless otherwise noted, all simulations were performed using GROMACS 2016 [67] and the Martini 2.2 force field [64]. The Berendsen thermostat was set to 313 K and a semi-isotropic Berendsen barostat was set to 1 bar in both directions (unless otherwise noted, see sect. 2.2.2). The compressibility modulus in both dimensions was 3 × 10^−5^ bar^−1^ with time constant 3 ps. Electrostatics and van der Waals (vdW) were shifted with a cutoff r-value of 1.2 nm. All systems underwent 100,000 steps of steepest-descent energy minimization prior to MD simulation with a timestep of 0.025 ps.

#### 2.2.2. Backbone restraint comparisons

Comparisons of different combinations of restraint scheme and run-time parameters took place in a 100% POPC membrane with E protein embedded. In two simulations, protein backbone beads were restrained to their initial coordinates via harmonic potential with a force constant of 2,000 kJ mol^−1^ nm^−2^. We describe this style of restraint as “absolute position restraints”. We performed two different simulations: one with run-time parameter *refcoord_scaling* set to *all*, and one with the setting turned off (*refcoord_scaling no*). Both simulations were then simulated in the NPT ensemble. The *refcoord_scaling no* simulation used GROMACS version 2021.6 and the v-rescale thermostat, but all other settings remained the same.

We also performed two simulations with elastic network restraints instead of absolute position restraints. In our elastic networks, all protein backbone bead pairs within 1.4 nm of one another were bonded together with force constants of 5,000 kJ mol^−1^ nm^−2^ using the *Martinize* tool [64]. One of these simulations (NPT ensemble) had identical settings to those listed in Common Settings. The second elastic network simulation (NVT ensemble) used GROMACS version 2021.6, the v-rescale thermostat, and utilized restraints to prevent diffusion, rotation, and tilt of the protein. Restraints were parameterized using the Colvars Dashboard [68].

The absolute position restraint simulation with *refcoord_scaling all* ran for over 50 *µs*, with the membrane adopting a metastable conformation before 10 *µs*. The absolute position restraint simulation with *refcoord_scaling no* and the NPT elastic network simulation ran for at least 10 *µs*, which was more than sufficient for area per lipid (∑) to converge. The NVT elastic network simulation ran for 2 *µs*.

#### 2.2.3. Mismatch and saturation comparisons

Membranes of varying mismatch level and lipid tail saturation (see Table 1) were simulated with the E protein embedded. Protein backbone beads were restrained with an elastic network which bonded all backbone bead pairs within 1.4 nm of one another with force constants of 5,000 kJ mol^−1^ nm^−2^. Five replicas of each system were simulated in the NPT ensemble for 10 *µs*. We discarded the first half of each trajectory prior to analysis.

To determine the equilibrium leaflet thickness *t*_0_ and equilibrium area per lipid ∑_0_ for each lipid species, we performed separate 2 *µs* NPT simulations of membranes composed of each lipid species listed in Table 1, but without a protein embedded. We discarded the first half of each trajectory prior to analysis. Thickness and area per lipid calculations are detailed in section 2.3.2.

### 2.3. Analysis

#### 2.3.1. Determination of hydrophobic mismatch

To determine the degree of hydrophobic mismatch present in each simulation system, we first calculated the length of the hydrophobic surface present on the E protein. To identify this surface, we identified the contiguously hydrophobic regions [69] of the protein sequence using the *blobulator* webtool [47]. We input the SARS-CoV E protein Uniprot ID (P59637) with a hydropathy cutoff of 0.6 and a minimum blob size of 10. With these settings, the only contiguously hydrophobic region present in the protein extends from residues V17 to A36.

We then measured the distance from V17 to the bottom of the CTD using the bond tool in VMD [70] and estimated the accessible hydrophobic surface to be approximately 1 nm thick, as depicted in Figure 1C. We then determined hydrophobic mismatch Δ*t* = *t*_TMD_ − *t*_0_ and relative mismatch Δ*t/t*_0_, where *t*_TMD_ is the protein hydrophobic surface half-thickness (0.5 nm) and *t*_0_ is the bulk equilibrium leaflet thickness, as shown in Table 1.

#### 2.3.2. Trajectory Analysis

Instantaneous area per lipid (∑ ≡ 2*L*^2^*/N*_*M*_, where *L* is the box length in the *x* or *y* direction and *N*_*M*_ is the number of lipid molecules in the membrane) was monitored throughout all simulation trajectories to ensure convergence.

All measurements of membrane profiles (height, thickness, curvature, asymmetry, etc.) were conducted using our membrane deformation analysis tool *nougat* [71]. If a protein was present, trajectories were wrapped, centered, and aligned around the protein backbone beads center of mass. Groups of lipid beads corresponding to a defined surface were binned on a polar lattice. Bins sizes were 1 nm in length radially and *π/*15 radians azimuthally. Averages were then taken within each bin in order to construct a surface for each frame in the simulation trajectory. In all cases analysis was performed on the second half of the trajectory to allow for proper system equilibration. Two-dimensional time averages were calculated by averaging over all frames included in the analysis (see *Figure S1*). One-dimensional profiles were calculated by averaging over the *θ* dimension of the time averages. In all cases the zone of analysis extends radially from *r* = *R*^(1,2)^ to *r* = *L/*2. Consequently, lipids in the corners of the simulation box and any lipids that may penetrate the pore of the protein are excluded from analysis.

Height surfaces are defined by different sets of lipid beads, depending on the surface in question. Cytoplasmic and luminal leaflet height surfaces *𝓏*^(1)^ and *𝓏*^(2)^ are calculated using the first bead in the acyl chain adjacent to the glycerol backbone (Martini beads C1A and C1B for those lipids present in this study). These beads in particular were chosen as they correspond to the hydrophobic surface of the membrane. To facilitate comparison between simulation systems, we transformed *𝓏*^(1,2)^ by the height of the center of mass of the TMD helices (*𝓏*_com_).

The interface separating the two leaflets *𝓏*^(*m*)^ is defined as the binned height average of any lipid tail bead within 6Å of any lipid tail bead belonging to the opposing leaflet. This interface surface is used to measure individual leaflet thicknesses *t*^(1)^≡ *𝓏*^(1)^ − *𝓏*^(*m*)^ and *t*^(2)^ ≡ *𝓏*^(*m*)^ − *𝓏*^(2)^. The bilayer midplane *𝓏*^+^ is the average of *𝓏*^(1)^ and *𝓏*^(2)^. The difference between bilayer interface and midplane fields is defined to be the leaflet thickness asymmetry *ε* ≡ *𝓏*^(*m*)^ − *𝓏*^+^ = (*t*^(2)^ − *t*^(1)^)*/*2, depicted in Fig. 2.

**Figure 2.**
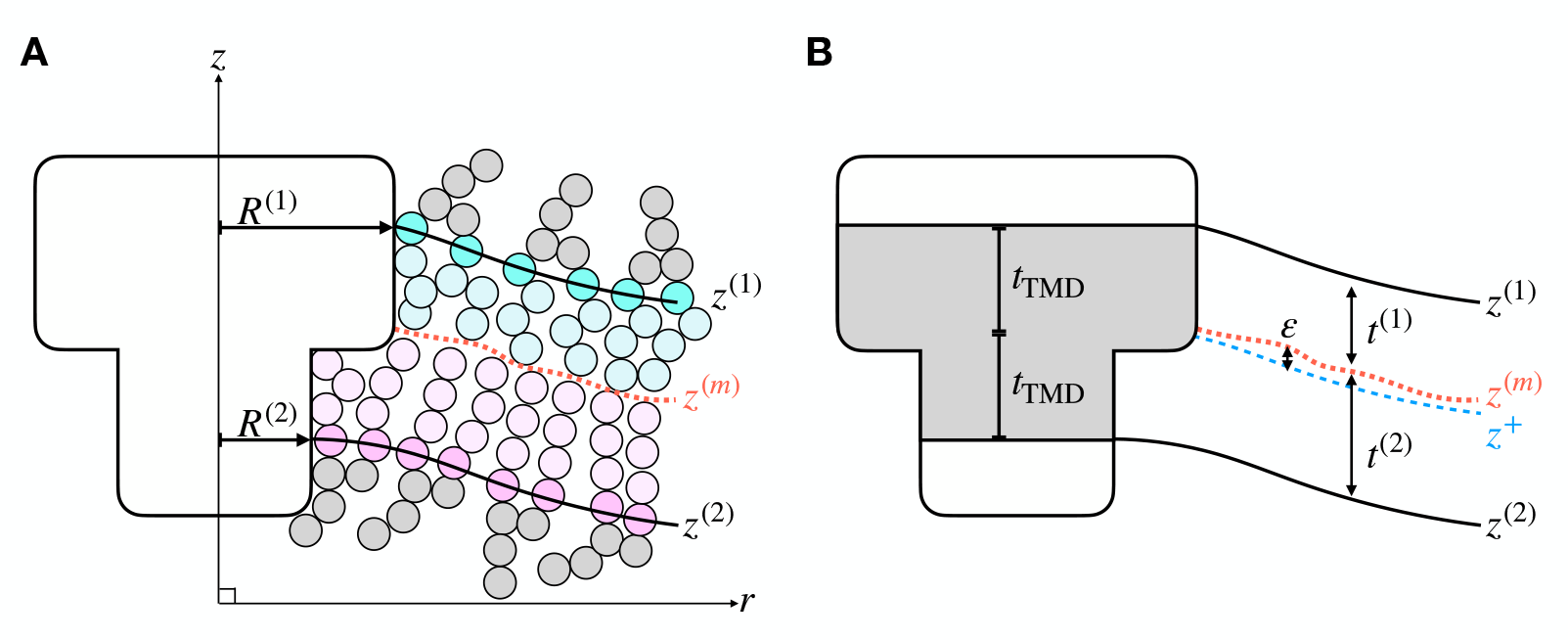
Cartoon depiction of lipid bilayer deforming around protein inclusion. *A)* A protein inclusion approximating the shape of the E protein pentamer with cytoplasmic leaflet radius *R*^(1)^ and luminal leaflet radius *R*^(2)^ is shown embedded in cartoon lipids forming a bilayer. Individual lipids are depicted with headgroups and glycerol backbones colored grey and acyl chains colored cyan (cytoplasmic leaflet) and pink (luminal leaflet). The outermost acyl chain beads are more saturated in color to indicate the hydrophobic surfaces *𝓏*^(1)^ and *𝓏*^(2)^. The leaflet interface surface *𝓏*^(*m*)^ is shown with a red dashed line. The fields *𝓏*^(1)^, *𝓏*^(2)^, and *𝓏*^(*m*)^ are measured using the membrane analysis package *nougat* [71]. Protein inclusion and lipids are not drawn to scale. *B)* The same inclusion and membrane as in *(A)*, but with cartoon lipids abstracted away. The thickness of the cytoplasmic leaflet (*t*^(1)^) and luminal leaflet (*t*^(2)^) is the distance in *𝓏* between *𝓏*^(*m*)^ and the hydrophobic leaflet surface (*𝓏*^(1)^ or *𝓏*^(2)^). The bilayer midplane *𝓏*^+^ (blue dashed line) is the average of *𝓏*^(1)^ and *𝓏*^(2)^. *𝓏*^+^ and *𝓏*^(*m*)^ differ due to local asymmetric leaflet thickness. The amount of local thickness asymmetry is defined as *ε* ≡ *𝓏*^(*m*)^ −*𝓏*^+^, and is proportional to the leaflet thickness difference. The inclusion’s hydrophobic region (grey shading) and its half-thickness (*t*_TMD_) are shown for educational purposes, but they do not match the hydrophobic region of the E protein, as shown in Figure 1C.

We assume the Monge gauge applies, where the membrane only deviates slightly from flat and the surface height can be expressed as a function of x and y. In this parameterization, mean curvature *H* is approximated as the negative Laplacian of the height surface (i.e. *H*^(1)^ ≡ − ∇^2^*𝓏*^(1)^ and *H*^(2)^ ≡−∇ ^2^𝓏 ^(2)^). We use the negative (as opposed to positive) Laplacian because we wish to follow the common convention that curvature should be positive if the osculating circle lies on the intracellular (luminal) side of the membrane.

In order to facilitate comparisons between our MD results and continuum theories of membrane deformation, we make use of a simplified version of the free energy expression in Ref. [63] eq. 18-20 with tilt deviations from the local surface normal assumed to be negligible 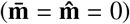, and the variable name for leaflet thickness substituted for consistency (*b* → *t*),

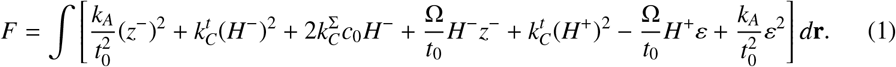

Here, as in Ref. [63], 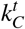 is the un-renormalized bending modulus that reflects the cost of bending the membrane while holding leaflet thickness equal to *t*_0_, 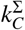 is the un-renormalized bending modulus that reflects the cost of bending the membrane while holding the interfacial area per molecule equal to its equilibrium area per lipid, *c*_0_ is the spontaneous curvature, *k*_*A*_ is the compressibility modulus, and Ω is a cross-term modulus. The field *𝓏*^−^ defines the thickness fluctuations of the membrane 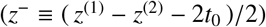 and *H*^−^ is the second derivative of *𝓏*^−^. We note that in the penultimate term of this expression *ε* and *H*^+^ are coupled, which we explore in more depth in section 3.5.

## 3. Results

### 3.1. Large-scale membrane bending is observed only under unintended compression

We aimed to reproduce the large-scale membrane bending shown in Ref. [48] by simulating the E protein pentamer in a 100% POPC membrane while varying the type of tertiary restraint used. When absolute position restraints were used to restrain the protein backbone beads to their initial coordinates via a harmonic potential, the membrane does exhibit long-range curvature (Fig. 3 A&B). With these settings, the membrane initially adopts a saddle configuration, with curvature propagating all the way to the edge of the simulation box (panel A). The saddle shape is stable for approximately 9 *µs*, at which point the raised edges of the membrane shift downward into a cylindrical conformation (panel B). This cylindrical conformation is stable for the rest of the trajectory (over 50 *µs*, see *Figure S2*). In contrast, when an elastic network is used to restrain nearby backbone beads to each other (Fig. 3 C&D), the membrane is globally flat with only slight deformation immediately local to the protein. This conformation persists throughout the simulation. These results showed that the membrane response to the E protein is particularly sensitive to the restraint and barostat settings, suggesting a potential explanation for large scale curvature observed in some studies.

**Figure 3.**
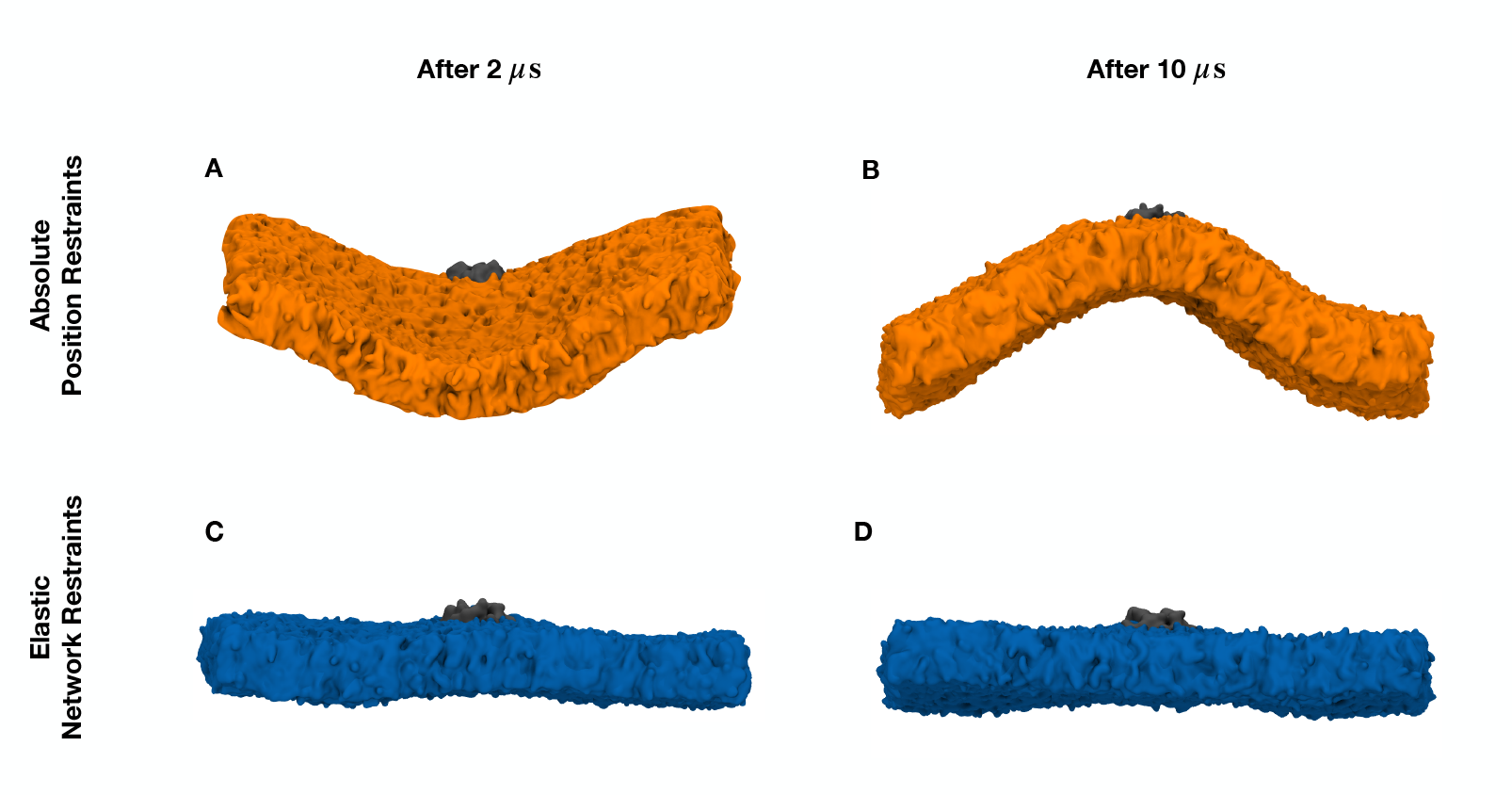
Membrane response to different protein restraint conditions. 100% POPC membrane with E protein embedded. Protein tertiary structure is restrained to initial coordinates via harmonic potential (“absolute position restraints”) *(A-B)* or nearby backbone beads are bonded to each other (“elastic network restraints”) *(C-D)*. All other settings and initial conditions are identical. Representative frames of the membrane conformation are taken after 2 *µs (A&C)* and 10 *µs (B&D)* of simulation.

We hypothesized that the difference in restraint settings might be influencing the projected area per lipid; if the membrane was compressed in plane, then out of plane bending would allow lipids to recover their equilibrium area. We consequently modulated simulation run-time parameters while monitoring the instantaneous area per lipid (∑) over the simulation trajectory. One such parameter, *refcoord_scaling*, exists for use with fixed restraints such as absolute position restraints. Its purpose is to temporarily scale the restraint coordinates with the rest of the simulation system when a barostat scales the system coordinates, speeding up system equilibration. While there are two options to choose from, *com* or *all*, GROMACS documentation notes that only *refcoord_scaling*=*all* will preserve the pressure and virial definitions when fixed restraints are used (see the Restraints documentation page in Ref. [72], accessed July 18, 2025). As shown in Fig. 4A, when *refcoord_scaling*=*all* is used in conjunction with absolute position restraints (orange line) the projected membrane area does not equilibrate; ∑ remains near its initial value of 0.6 *nm*^2^ for the duration of the trajectory. In contrast, turning off *refcoord_scaling* (*refcoord_scaling*=*no*, red line) allows the membrane to equilibrate normally and ∑ quickly reaches the reference value for POPC ( ≈ 0.67 *nm*^2^). This mirrors the membrane behavior seen in our elastic network simulation (blue line), which did not use *refcoord_scaling*. As can be seen in Fig. 4B, *refcoord_scaling*=*no* (red membrane) also yields the same nearly-flat membrane as when an elastic network is used (blue membrane). In summary, the use of *refcoord_scaling*=*all*, in conjunction with absolute position restraints, appears to prevent box scaling in this system.

**Figure 4.**
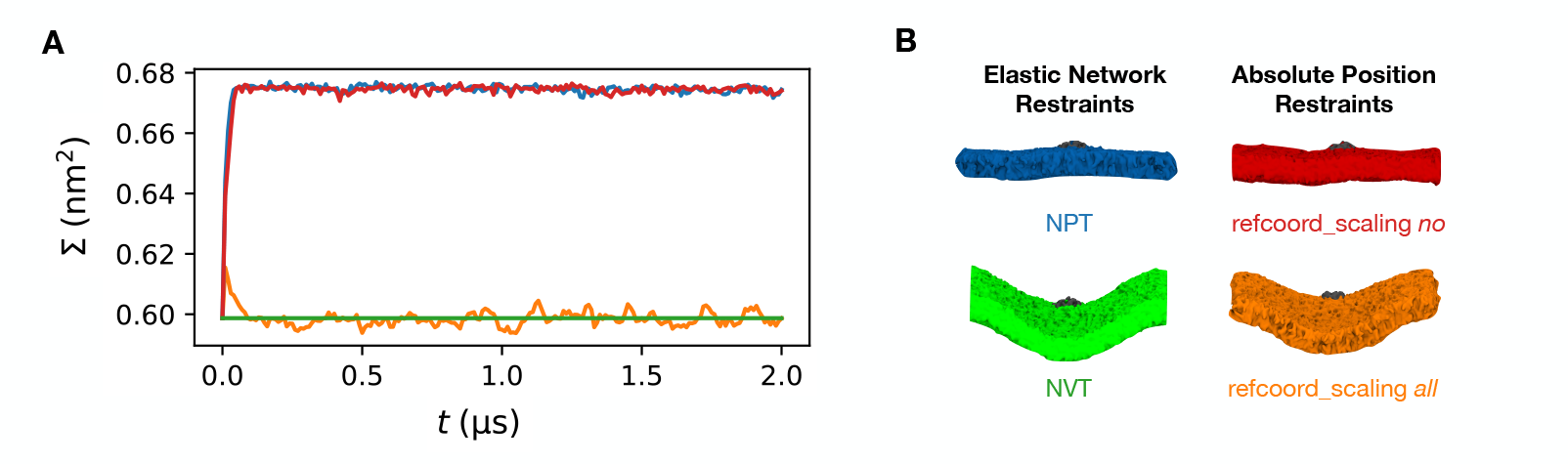
Membrane bending is dependent upon pressure scaling settings. *A)* Instantaneous area per lipid ∑ ≡ 2*L*^2^*/N*_*M*_ for a 100% POPC system utilizing four different combinations of restraint scheme and GROMACS barostat run-time parameter: elastic network restraints and constant pressure (NPT) simulation (blue), elastic network restraints and constant volume (NVT) simulation (green), absolute position restraints and *refcoord_scaling no* (red), absolute position restraints and *refcoord_scaling all* (orange). *B)* Representative frames of the systems shown in *A*, taken after 2 *µs* of simulation. Lipids colored by restraint scheme, as in *A*.

To test whether the membrane bending observed above was solely a product of the membrane being unable to reach its equilibrium area per lipid while remaining flat, we performed a simulation using an elastic network and no barostat (NVT). Both ∑ (green line, Fig. 4A) and membrane deformation (green membrane, Fig. 4B) match our results from the *refcoord_scaling*=*all* simulation (orange line/membrane), which is consistent with membrane bending induced by compression.

Taken together, this evidence indicates that the combination of absolute position restraints and *refcoord_scaling*=*all* kinetically traps the simulation box. Unless the simulation begins with a lipid area density that is very close to the equilibrium value, the membrane bends to reduce compression of the individual lipids. We believe this scenario may account for the curvature reported by Mehregan et al. [48]. More significantly, this outcome may be relevant to other anisotropic coarse-grained MD simulations involving fixed restraints. This outcome may be relevant to other anisotropic coarse-grained MD simulations involving fixed restraints. We were able to properly equilibrate these membranes by using an elastic network in lieu of absolute position restraints, or using refcoord_scaling=com with absolute position restraints. We observed this behavior across several versions of GROMACS. We have not yet investigated whether this effect would significantly affect simulations that do not contain box-spanning membranes. We also note that this scenario is distinct from the neighbor list artifact detected in GROMACS by Kim et al. [73].

### 3.2. E protein does not induce long-range curvature in uncompressed membranes

To determine whether the near-flat behavior of the POPC membrane shown above (Fig. 3 C&D) generalizes to membranes of varying mismatch and lipid saturation, we simulated the E protein embedded in 9 homogeneous membranes (see table 1) and measured the average height difference of each leaflet (*𝓏*^(1,2)^ − *𝓏*_com_, where *𝓏*_com_ is the height of the center of mass of the E protein TMD helices, allowing for comparison between simulation systems) as a function of distance *r* from the protein center (Fig. 5 A&B). To ensure that the barostat behavior described above did not affect our results, we utilized elastic network restraints in all simulations. In all cases, the height of the leaflet surface at the protein-leaflet interface (*r* = *R*) is slightly higher than the height of the leaflet surface at *r* = *L/*2, indicating that the membrane bends “downwards” and away from the protein CTD. All cytoplasmic leaflets (solid lines) are comparatively flat, exhibiting a total height change of less than one nanometer. Deformation is more pronounced in the luminal leaflet (*𝓏*^(2)^, dashed lines), where the height decrease is sharpest within the first few lipidation shells before an inflection point is reached (*r* ≈ 3.5 nm) and the decrease becomes much more gradual with *r*. As shown in Fig. 5C the distance required to halve luminal leaflet height (*r*_50_) decreases with mismatch, demonstrating that increasing the mismatch level increases the severity of the local luminal leaflet deformations. In summary, while we observe severe local deformations, there is only slight long-range membrane bending even under extreme mismatch conditions. This bending does not appear to achieve the scale necessary to be consistent with the E protein’s purported role in driving the budding process, although multiple E proteins and/or other membrane proteins may interact to achieve this effect.

**Figure 5.**
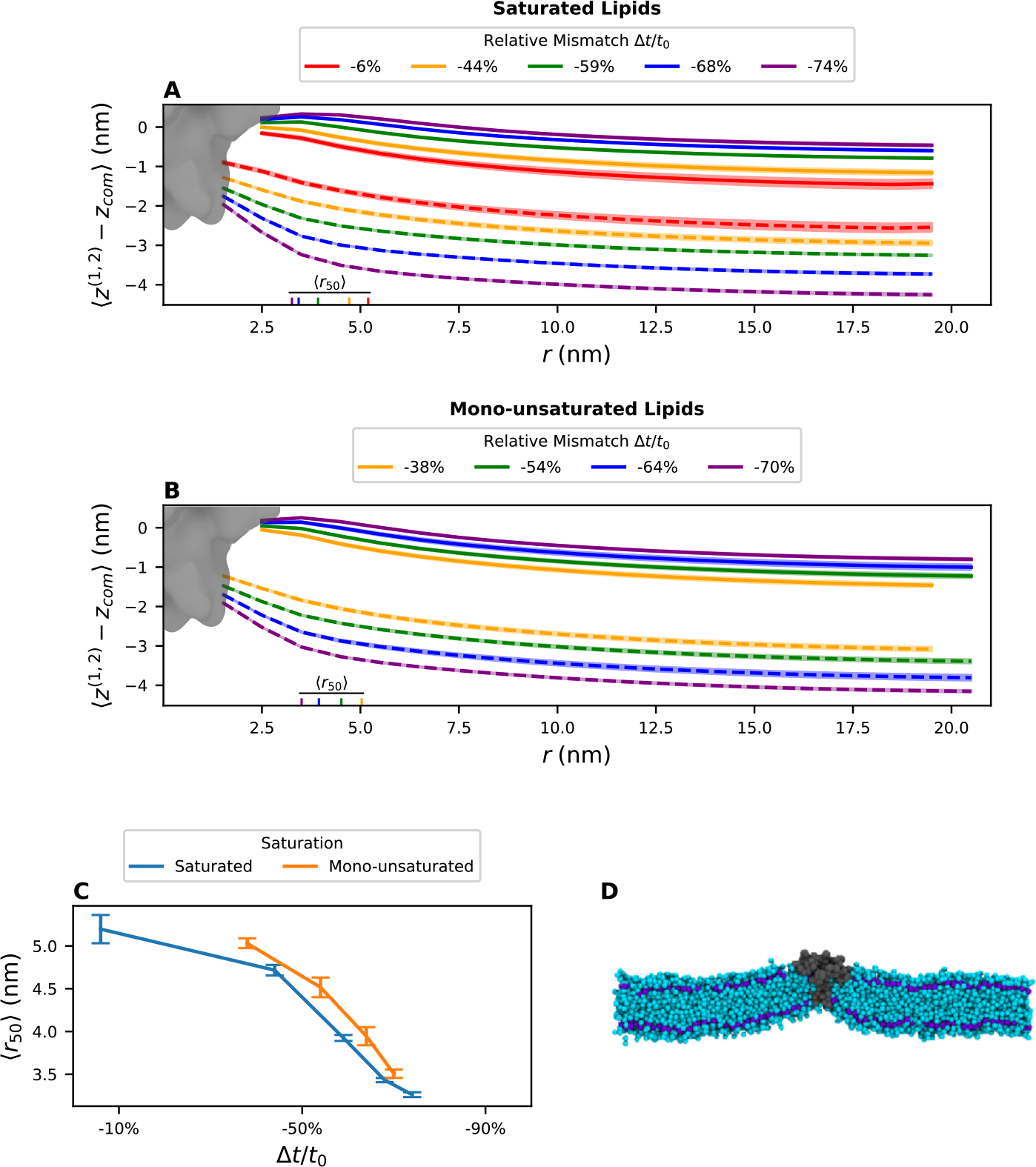
One dimensional height profiles of membranes with varying degrees of mismatch and saturation. Average height difference *𝓏*^(1,2)^ − *𝓏*_com_, where *𝓏*_com_ is the center of mass of the protein TMD helices, as a function of distance from the protein center *r* for saturated *(A)* and mono-unsaturated *(B)* lipid membranes with negative hydrophobic mismatch values. Solid lines denote cytoplasmic leaflet surfaces while dashed lines denote luminal leaflet surfaces for each membrane. The approximate position of the E protein is shown in grey. *r*_50_, the distance at which the luminal leaflet surface has attained half its total decrease in height from *r* = *R*^(2)^ to *r* = *r*_max_, is denoted along the *r* axis. All averages are taken over the second half of a 10 *µs* simulation, over 5 replicas. 95% confidence interval indicated with shading. *C) r*_50_ for saturated (blue) and mono-unsaturated (orange) lipid membranes, as a function of degree of mismatch Δ*t/t*_0_. Error bars represent 95% confidence interval across five replicas. *D)* Representative frame of saturated membrane with Δ*t/t*_0_ = 74%, captured after 10 *µs* of simulation and then cross-sectioned to reveal membrane interior. Lipids are depicted in cyan and the E protein in gray. The lipid beads comprising the hydrophobic surfaces *𝓏*^(1,2)^ are depicted in purple. Water and ions are present in the simulation system, but hidden for clarity.

### 3.3. Hydrophobic mismatch contributes to local membrane deformations

To investigate how changing the level of hydrophobic mismatch and lipid flexibility affects the thickness and curvature profiles of the membrane, we measured normalized membrane thickness and membrane mean curvature as a function of *r* (Fig. 6A). Membrane thickness normalized by its bulk equilibrium thickness 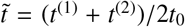 (Fig. 6A, top) shows that the greatest change in deformation occurs immediately local to the protein. At *r* = *R*, all membranes have 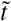 minima significantly less than 100% of equilibrium thickness, indicative of membrane compres-sion. 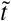 then “overshoots” its bulk equilibrium value, peaking around *r* ≈ 3.5 − 5.5 nm before decreasing to 100% in the bulk, regardless of mismatch level. In both saturated and monounsaturated lipid membranes, normalized thickness at the protein-membrane interface 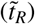 (Fig. 6B, top) decreases monotonically with degree of mismatch. Saturated membranes have lower 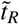 than mono-unsaturated membranes of the same equilibrium thickness, suggesting that the additional flexibility introduced by the double bond makes the membrane more accommodating of hydrophobic mismatch. The finding that thickness deformations only persist in the immediate vicinity of the protein and do not extend into the bulk is consistent with previous works that found membranes heal back to equilibrium thickness within a few lipidation shells, regardless of inclusion radius and degree of mismatch [74, 75]. The thickness overshoots, where a compressed membrane initially exceeds its equilibrium thickness before reaching unity, have been previously predicted in e.g. refs. [58, 59, 61, 76] and observed computationally in e.g. ref. [75].

**Figure 6.**
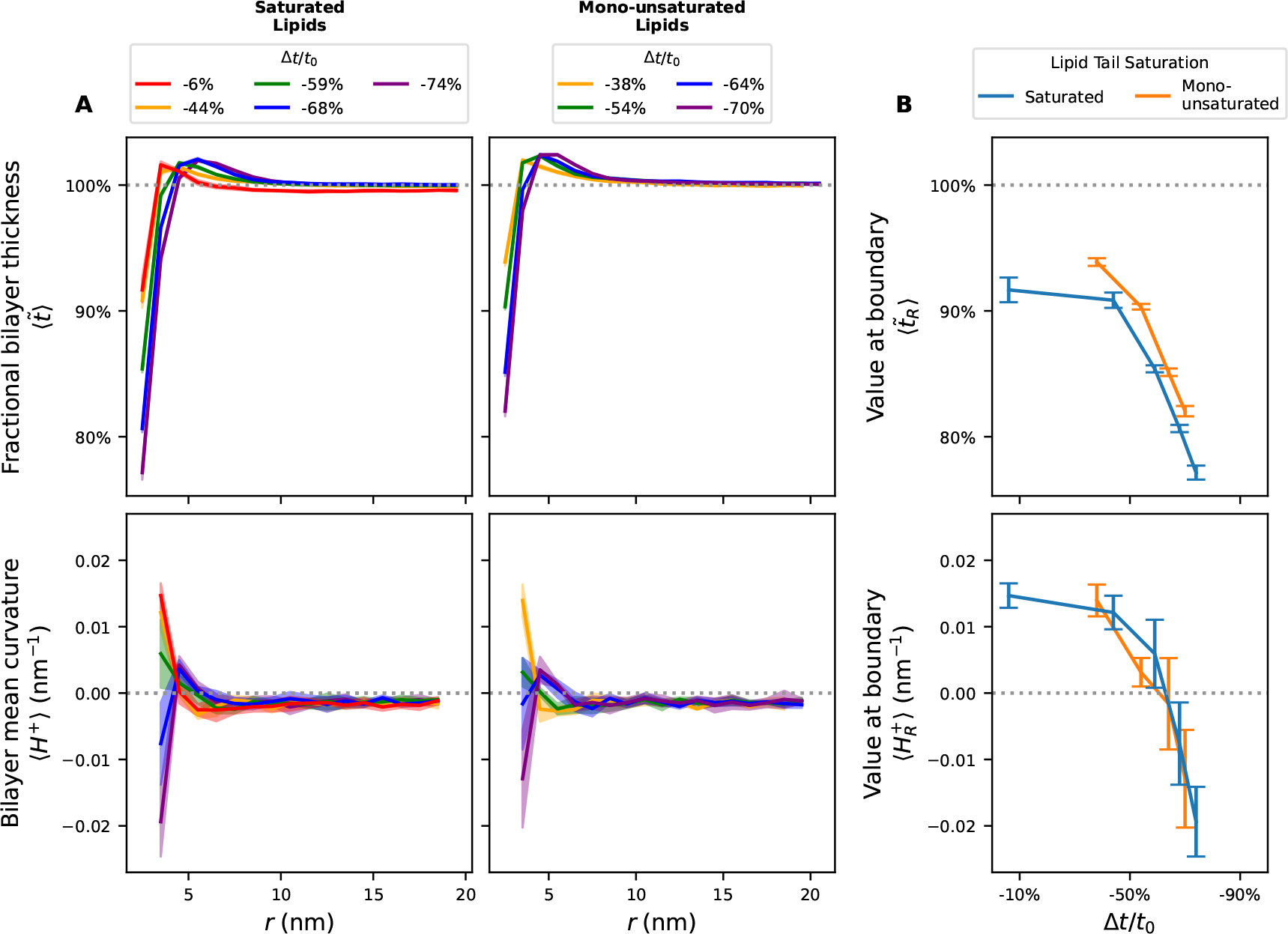
Membranes thin and bend as they approach the E protein. A) Fractional bilayer thickness 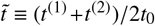 and bilayer mean curvature H^+^ ≡ ™∇^2^𝓏^+^ (bottom) as a function of distance r from the protein center for saturated(left) and mono-unsaturated (right) lipid bilayers with negative hydrophobic mismatch values. All averages taken overthe second half of a 10 μs simulation, across 5 replicas. 95% confidence interval indicated with shading. Dotted greylines are shown at 100% in thickness plots or 0 in mean curvature plots to guide the eye. B) Average 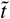 (top) and H+(bottom) values at the protein-membrane interface r = R^(1)^ across the 5 replicas. Error bars represent 95% confidenceinterval.

To further isolate membrane bending from the effect of increasing the equilibrium thickness of the membrane, we measured the bilayer mean curvature *H*^+^ as a function of *r* (Fig. 6A, bottom). As noted in section 2.3.2, we followed the convention that curvature should be positive if the osculating circle lies on the intracellular side of the membrane. In all cases the highest magnitude mean curvature is at *r*=R, with signed values ranging from positive to negative with the degree of mismatch. As *r* increases the curvature trends to near-zero in all cases, indicating that while there is slight curvature in the bulk, it is at least an order of magnitude lower than that near the protein. Furthermore, bulk (*r* > ≈ 7.5 nm) curvature does not differ with the degree of mismatch, confirming that the slight bending observed in Fig. 5 is unrelated to hydrophobic mismatch. Protein-membrane interface curvature values 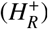 for saturated and mono-unsaturated lipid membranes are within error of each other (Fig. 6B, bottom), indicating that membrane flexibility does not alter the curvature profiles significantly.

### 3.4. Individual leaflet deformations are asymmetric

While thickness differences between leaflets have previously been assumed to be negligible in the case of cylindrically symmetric inclusions (e.g. in Ref. [75]), the non-cylindrical shape of the E protein and the presence of the leaflet thickness asymmetry term in eq. 1 caused us to revisit this assumption. The leaflet thickness asymmetry *ε* is proportional to the thickness difference between the two membrane leaflets. To test whether asymmetry is negligible in the case of this protein, we measured relative leaflet asymmetry *ε/t*_0_ as a function of *r*. Positive values correspond to a cytoplasmic leaflet that is thinner than the luminal leaflet. As shown in Figure 7A (top panels), *ε/t*_0_ is positive everywhere except for the thickest mono-unsaturated membrane. Asymmetry at the protein-membrane interface increases with degree of mismatch, ranging from 1% to 9% of equilibrium thickness. In all cases but one, *ε/t*_0_ converges to a positive, non-zero value at *r >* 5 nm, suggesting that not only is asymmetry significant local to the protein, it is also non-vanishing in the bulk when curvature is present. There do not appear to be significant differences in asymmetry depending on lipid saturation (Fig 7B) with the possible exception of the mono-unsaturated lipid membrane with Δ*t/t*_0_ =− 64%.

**Figure 7.**
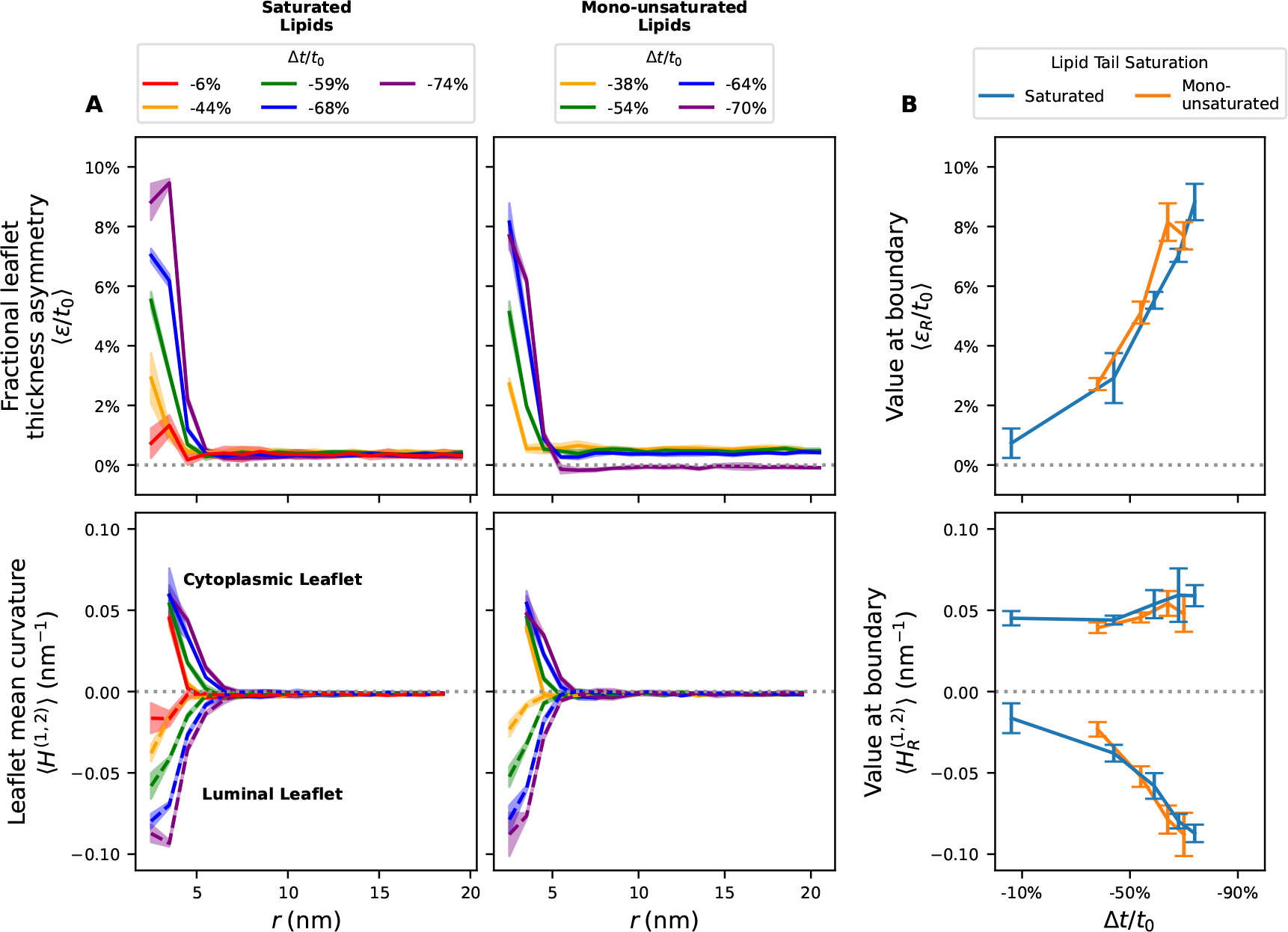
E protein deforms individual leaflets asymmetrically. *A)* Fractional leaflet thickness asymmetry *ε/t*_0_ (top) and leaflet mean curvature *H*^(1,2)^ (bottom) as a function of distance *r* from protein center for saturated (left) and monounsaturated (right) lipid bilayers with negative hydrophobic mismatch values. Solid lines denote cytoplasmic leaflet curvature *H*^(1)^ and dashed lines denote luminal leaflet curvature *H*^(2)^ (bottom). All averages taken over the second half of a 10 *µs* simulation, across 5 replicas. 95% confidence interval indicated with shading. Dotted grey lines are shown at 0 to guide the eye. *B)* Average values of *ε* (top) and *H*^(1,2)^ at protein-membrane interface *r* = *R*^(1)^ (*ε*_*R*_, *H*^(1)^) or *r* = *R*^(2)^ (*H*^(2)^) across the 5 replicas. Error bars represent 95% confidence intervals.

To determine whether individual leaflet curvature is also asymmetric, we measured leaflet mean curvature *H*^(1,2)^ as a function of *r* (Fig. 7A, bottom). Cytoplasmic leaflet surfaces have positive curvature (solid lines) and achieve their global maximum at the leaflet-protein interface (*r*=R) before decreasing monotonically with *r* towards zero. Luminal leaflet surfaces have negative curvature (dashed lines) and in all but the most mismatched membrane achieve a global minimum at the protein interface and then increase monotonically with *r* toward zero curvature. As expected, the magnitude of the curvature at the protein-leaflet interfaces 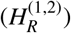 increases with degree of mismatch (Fig. 7B, bottom). The curvatures are not symmetric across zero for the two leaflets, which may be expected due to the differing boundary conditions of the luminal and cytoplasmic leaflets due to the interactions of the E protein CTD with the cytoplasmic leaflet.

### 3.5. Curvature and leaflet thickness asymmetry coupling helps minimize free energy

To verify whether or not the combination of mean curvature *H*^+^ and leaflet thickness asymmetry *ε* produces a negative free energy term, we measured −*εH*^+^ directly (Fig. 8 A, top panels). Notably, all systems average to negative values for all *r*. At the protein-membrane interface 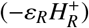 this effect increases with mismatch and when lipids are mono-unsaturated (Fig. 8B, top). Provided that the cross-term modulus Ω (see Eq. 1) is a positive constant, the combination of curvature and leaflet thickness asymmetry creates a negative term that might stabilize an otherwise costly membrane conformation.

**Figure 8.**
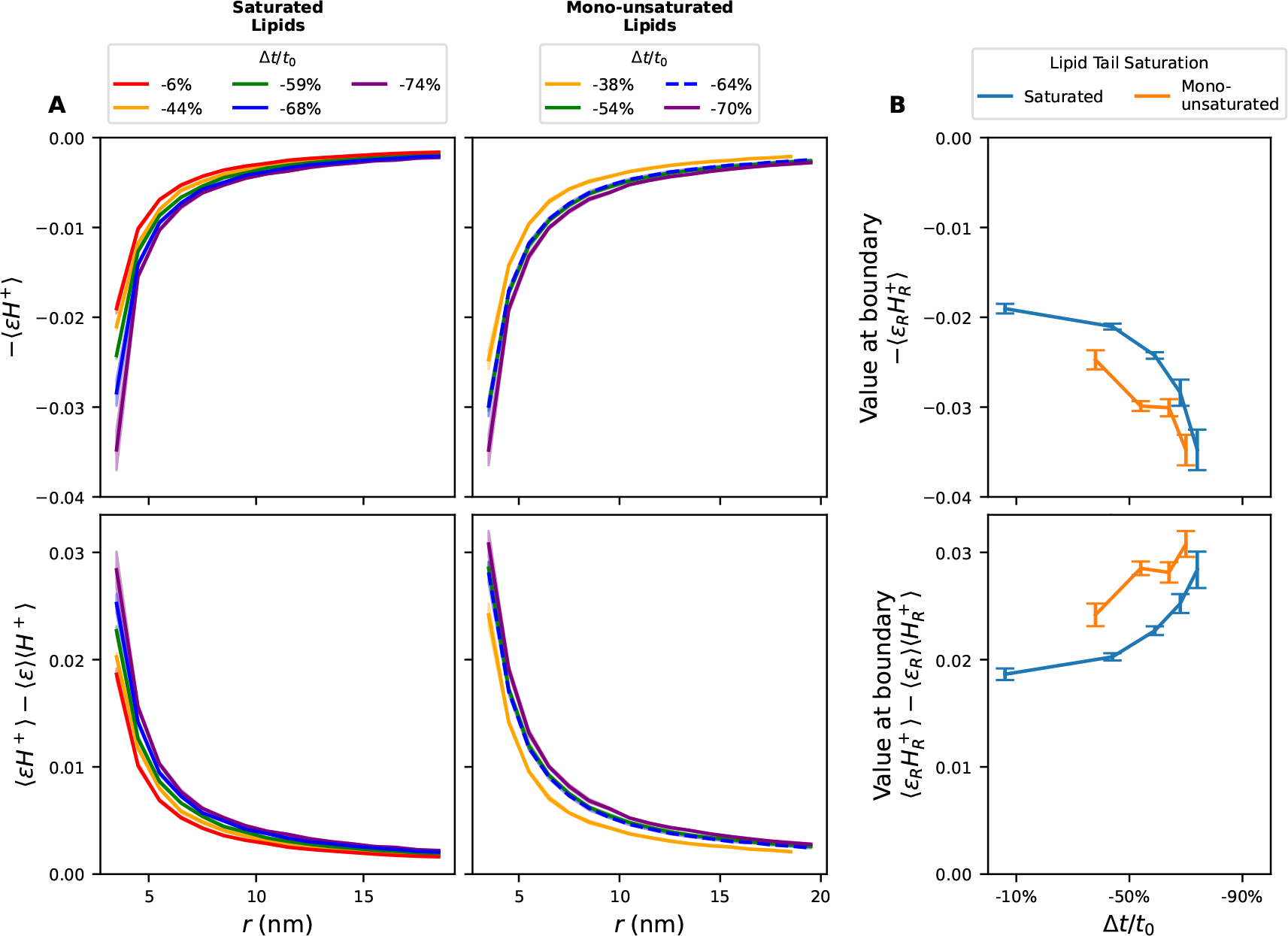
Thickness asymmetry and mean curvature produce a negative energy term and are correlated. *A)* The negative product of leaflet thickness asymmetry *ε* and bilayer mean curvature *H*^+^ (top) and the correlation of the two ( ⟨*εH*^+^⟩ −⟨*ε*⟩⟨*H*^+^⟩, bottom) as a function of distance from the protein center, *r*, for saturated (left) and mono-unsaturated (right) lipid bilayers with negative hydrophobic mismatch values. All averages taken over the second half of a 10 *µs* simulation. 95% confidence interval indicated with shading. The mono-unsaturated lipid bilayer with -54% mismatch (green line) is present in the right panels of A, but obscured by the bilayer with -64% mismatch (dashed blue line). *B)* Average values of −*εH*^+^ (top) and ⟨*εH*^+^⟩ − ⟨*ε*⟩⟨*H*^+^⟩ (bottom) at protein-membrane interface *r* = *R*^(1)^ across the 5 replicas. Error bars represent 95% confidence intervals.

As *ε* and *H*^+^ are coupled in the simplified bending free energy expression (eq. 1), we calculated the correlation ⟨*εH*^+^⟩− ⟨*ε*⟩ ⟨*H*^+^⟩ as a function of distance *r* as shown in Fig. 8A (bottom panels). Positive correlation indicates that when curvature is positive, the cytoplasmic leaflet is thinner on average than the luminal leaflet. Correlation is highest at *r*=R, decreases with *r*, and is always positive. Correlation increases with the degree of mismatch, and is higher for more flexible lipids (Fig. 8B, bottom). These results show that curvature and thickness deformations are positively correlated in the bulk, but are significantly more correlated adjacent to the E protein. This indicates that while coupling between the terms may be negligible for bulk membrane fluctuations, it becomes significant around inclusion boundaries. In particular, this may affect inclusions with complex surfaces like the E protein.

## 4. Conclusion

Our results suggest that, in isolation, the E protein of SARS-CoVs 1 and 2 does not cause long-range membrane bending consistent with a role in budding. These results corroborate the findings of Collins et al. [49] and Kuzmin et al. [50], but use an experimentally-validated protein structure [25] rather than a predicted structure [77]. Additionally, we have found that the combination of absolute position restraints on protein conformation with a particular run-time parameter (*refcoord_scaling all* in GROMACS) can can lead to a kinetically trapped membrane state under certain barostat settings. We believe this trapped state particularly affects coarsegrained membrane simulations that use fixed restraints, and that this may explain the long-range membrane curvature observed previously in simulation [48].

While long-range bending is minimal, short-range membrane and leaflet deformations immediately surrounding the protein are notable, even in the case of the lipids with only 8-carbons per acyl chain (DTPC Δ*t/t*_0_ = − 6%, *t*_0_ = 0.53 nm). In all cases the leaflet thickness asymmetry *ε/t*_0_ is non-zero near the protein-membrane interface. As membrane relative mismatch grows, local deformations to membrane and leaflets are significantly exacerbated. Consequently, asymmetric hydrophobic mismatch is established as a primary driver of asymmetric membrane deformation around non-cylindrical inclusions. Further work is required to disambiguate asymmetric hydrophobic mismatch conditions from those conditions arising from wedge-shaped inclusions classically considered under the contact angle mechanism framework [53].

Varying the stiffness of the membrane via saturation level also offered insight into the difference in membrane reaction to an irregular inclusion like the E protein. The less stiff mono-unsaturated lipid membranes were less compressed, and exhibited stronger coupling between leaflet thickness asymmetry and leaflet mean curvature than saturated lipid membranes at the same or similar levels of mismatch. It is unclear whether this trend will hold if more flexible lipids than those used here are introduced, but the introduction of leaflet thickness asymmetry appears to be a viable and non-negligible way for membranes to alleviate inclusion-induced deformation.

## Supporting information

Custom lipid .itp file

## Acknowledgments

The authors would like to acknowledge conversations with Frank L.H. Brown. This material is based upon work supported by the National Science Foundation under Grant No. DGE 2152059. Computational resources were provided by the Rutgers Office of Advanced Research Computing.

## Data Availability

Three Zenodo repositories have been published containing all simulation trajectories and analysis files used in this paper. An additional repository contains a minimal working example demonstrating the barostat behavior identified in this paper. They can be found at the following addresses:

- Restraint comparison simulations: https://doi.org/10.5281/zenodo.15238548
- Minimal example of E protein in smaller POPC membrane: https://doi.org/10.5281/zenodo.16109228
- Saturated lipid simulations: https://doi.org/10.5281/zenodo.16106026
- Mono-unsaturated lipid simulations: https://doi.org/10.5281/zenodo.16110026

## Notes

### Competing Interest Statement

The authors have declared no competing interest.

### Summary of Updates

Some sections were rewritten for clarity; some language was softened; symbols that were only used once were dropped from the definitions list and defined locally instead.

