## Supplementary figures for "SARS-CoV E protein couples asymmetric leaflet thickness and curvature deformations"

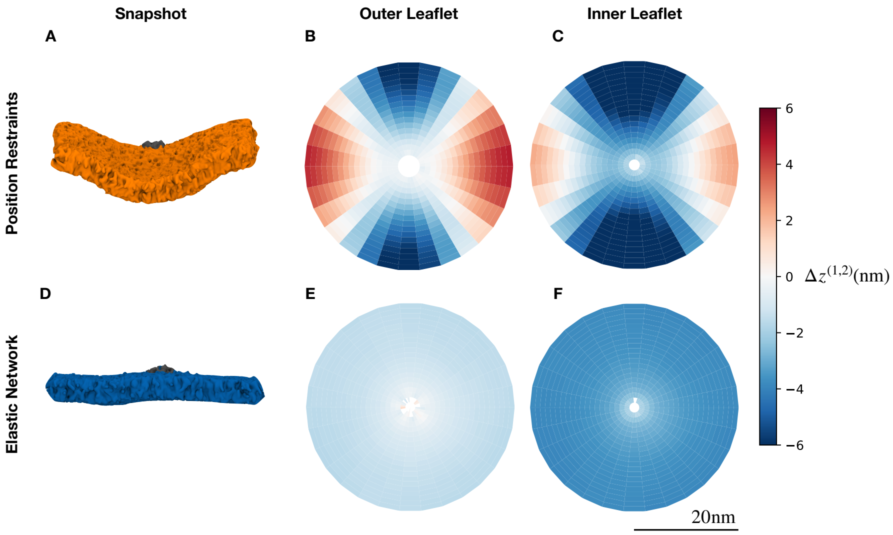


Figure S1: **Illustration of nougat leaflet height measurement.** Still images *(A, D)* and 2D height profiles of cytoplasmic *(B, E)* and luminal *(C, F)* leaflets from two simulation systems, one with absolute position restraints restraining the protein backbone beads to their initial coordinates in space *(A-C)*, and one with an elastic network restraining the protein backbone beads to each other *(D-F)*. Heatmaps *(B-C, E-F)* display the leaflet height ($\Delta z^{(1,2)} \equiv z^{(1,2)} - z_{com}$, where $z_{com}$ is the height of the protein TMD's center of mass), averaged over the trajectory and presented as a function of $r$ and $\theta$. The protein's center of mass is centered at the origin. 1D profiles can be generated (as seen in the paper) by averaging over the $\theta$ dimension.

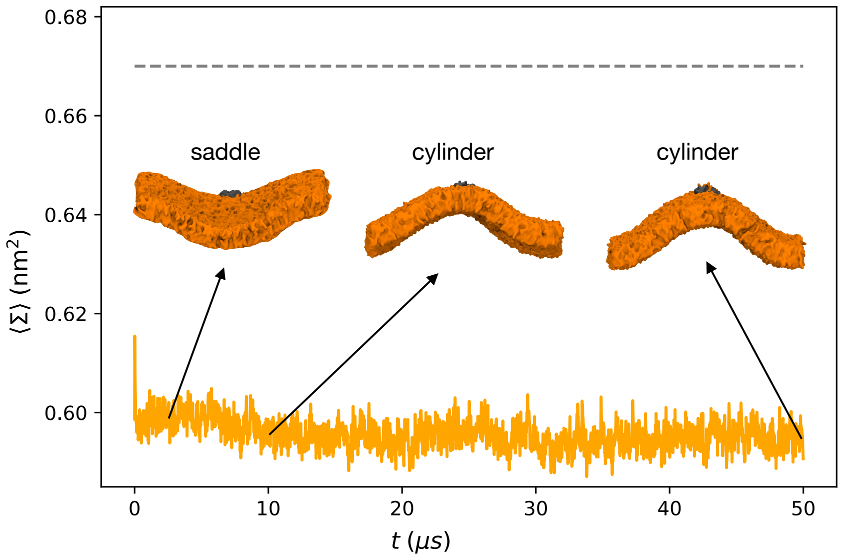


Figure S2: **Instantaneous area per lipid** (Σ) of the absolute position restraint system with *refcoord_scaling=all*, monitored over the course of the full 50 $\mu s$ trajectory. Snapshots are shown, taken at 2, 10, and 50 $\mu s$ respectively, along with their approximate shape annotated in text. The dashed grey line indicates the equilibrium area per lipid for POPC, 0.67 $nm^2$.

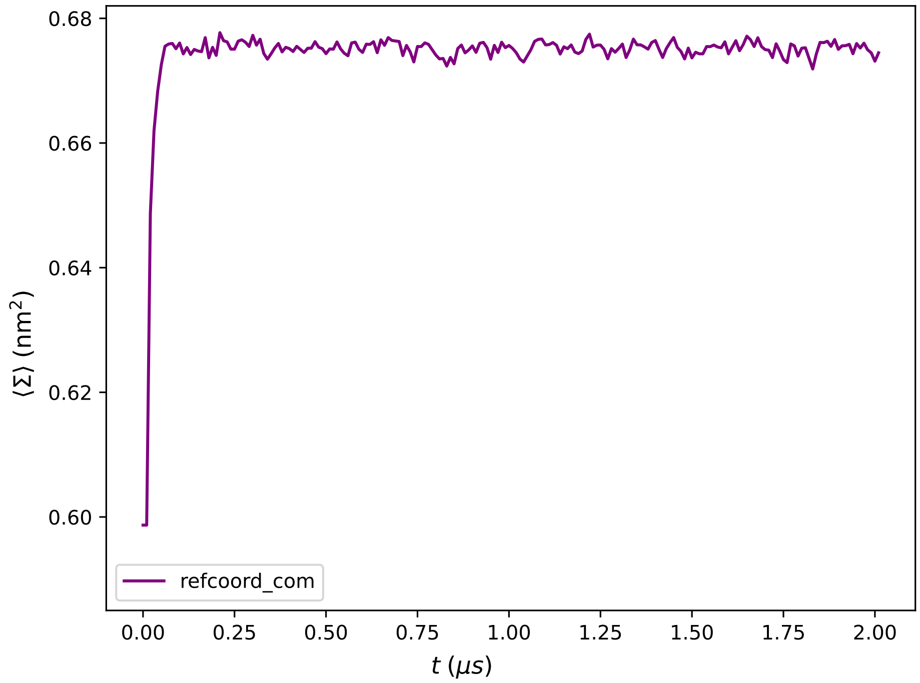


Figure S3: **Instantaneous area per lipid** $(\Sigma)$ for a 100% POPC system that uses absolute position restraints and *refcoord_scaling=com*.

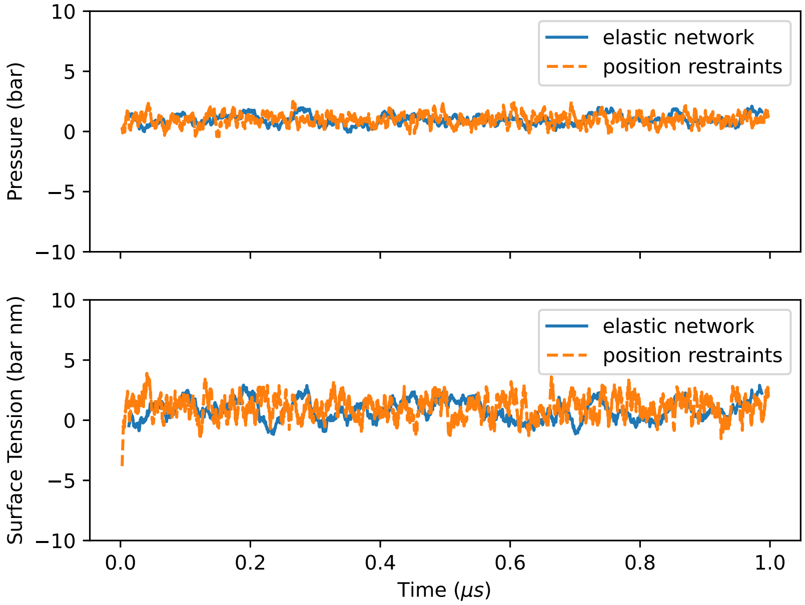


Figure S4: **Comparison of diagnostic values.** Pressure (top) and surface tension (bottom) from first $\mu s$ of elastic network simulation (blue) and absolute position restraints (orange) simulation systems. Values captured by *gmx energy*, then smoothed with rolling average window width of 25. Values at time 0 are omitted.
